# Structural and functional plasticity in the dorsolateral geniculate nucleus of mice following bilateral enucleation

**DOI:** 10.1101/2020.11.02.365130

**Authors:** Ashish Bhandari, Thomas W. Ward, Jennie Smith, Matthew J. Van Hook

## Abstract

Within the nervous system, plasticity mechanisms attempt to stabilize network activity following disruption by injury, disease, or degeneration. Optic nerve injury and age-related diseases can induce homeostatic-like responses in adulthood. We tested this possibility in the thalamocortical (TC) neurons in the dorsolateral geniculate nucleus (dLGN) using patch-clamp electrophysiology, optogenetics, immunostaining, and single-cell dendritic analysis following loss of visual input via bilateral enucleation. We observed progressive loss of vGlut2-positive retinal terminals in the dLGN indicating degeneration post-enucleation that was coincident with changes in microglial morphology indicative of microglial activation. Consistent with the decline of vGlut2 puncta, we also observed loss of retinogeniculate (RG) synaptic function assessed using optogenetic activation of RG axons while performing whole-cell voltage clamp recordings from TC neurons in brain slices. Surprisingly, we did not detect any significant changes in the frequency of miniature post-synaptic currents (mEPSCs) or corticothalamic feedback synapses. Analysis of TC neuron dendritic structure from single-cell dye fills revealed a gradual loss of dendrites proximal to the soma, where TC neurons receive the bulk of RG inputs. Finally, analysis of action potential firing demonstrated that TC neurons have increased excitability following enucleation, firing more action potentials in response to depolarizing current injections. Our findings show that degeneration of the retinal axons/optic nerve and loss of RG synaptic inputs induces structural and functional changes in TC neurons, consistent with neuronal attempts at compensatory plasticity in the dLGN.

## Introduction

The activity of neuronal networks in the brain is one of the most stable components of the nervous system and this stability is required throughout life to maintain functional connectivity for effective information transfer (Turrigiano, 1999, 2008; Turrigiano and Nelson, 2004; D’Angelo, 2010). This results from the nervous system’s ability to monitor changes in network activity and compensate via modulation of neuronal structure, intrinsic excitability, and/or synaptic function (Abbott and Nelson, 2000; Darlington et al., 2002; Schulz, 2006; Beck and Yaari, 2008; Howland and Wang, 2008; Gainey et al., 2009; Turrigiano, 2012; Lambo and Turrigiano, 2013; Bliss et al., 2014; Hauser et al., 2014; Fernandes and Carvalho, 2016; Chowdhury and Hell, 2018; Meriney and Fanselow, 2019). Such plasticity in cortical and sub-cortical structures has been reported during maturation as well as in adulthood in response to learning and memory or injury and is an important feature of the nervous system during learning and development as well as in disease.

One of the key structures of the visual system is the dorsolateral geniculate nucleus (dLGN), which receives direct projections from retinal ganglion cells, integrates visual responses (Mazade and Alonso, 2017; Monavarfeshani et al., 2017), and relays that information to the primary visual cortex (Sherman and Guillery, 2002; Chen et al., 2016). Disruption in the flow of information resulting from injury or disease will interrupt normal communication and may trigger compensatory mechanisms to counter the disruption. The developmental plasticity of the dLGN is well-established and involves pruning and strengthening of synaptic inputs from retinal ganglion cells, refinement of dLGN relay neuron dendritic branching, and maturation of intrinsic excitability (Chen and Regehr, 2000; Hooks and Chen, 2006, 2008; Bickford et al., 2010; Hong and Chen, 2011; Krahe et al., 2012; El-Danaf et al., 2015; Thompson et al., 2016; Litvina and Chen, 2017a; Guido, 2018). A handful of studies have also documented plasticity in the dLGN in adult mice, including in monocular deprivation (Jaepel et al., 2017; Rose and Bonhoeffer, 2018), monocular enucleation (Gonzalez et al., 2005), a mouse model of inflammatory demyelination (Araújo et al., 2017), and ocular hypertension (Bhandari et al., 2019; Van Hook et al., 2021). These studies have shown that alterations in visual function or optic nerve health lead to changes in inputs to the dLGN from retina and the cortex as well as changes in the structure and function of dLGN relay neurons (Yücel et al., 2001; Hayakawa and Kawasaki, 2010; Krahe and Guido, 2011; Ly et al., 2011; Krahe et al., 2012; You et al., 2012; El-Danaf et al., 2015; Pang et al., 2015; Araújo et al., 2017; Bhandari et al., 2019).

Given that plasticity in the dLGN has been well established during development as well as adulthood, it is still unknown how the mature dLGN responds to severe optic nerve injury. While we have previously explored the influence of ocular hypertension (Bhandari et al., 2019; Van Hook et al., 2021), which induces a low-level and chronic optic nerve injury and alteration in RGC output, our goal here was to use a more dramatic insult to probe plasticity processes in the dLGN. To accomplish this, we bilaterally enucleated adult mice to test the hypothesis that traumatic optic nerve injury will induce functional changes in the neurons and synapses in the dLGN.

### Experimental Procedures

#### Animals

All procedures were approved by the Institutional Animal Care and Use Committee at the University of Nebraska Medical Center. C57 Bl/6J mice (Jackson Labs #000664) were used for excitability experiments. For optogenetic experiments, Chx10-cre;Ai32 (Litvina and Chen, 2017b) mice were used, generated by crossing Chx10-cre (Rowan and Cepko, 2004) (Chx10 BAC Jackson labs #005105) mice with Ai32 (Madisen et al., 2012) reporter line (Jackson labs #024109). Chx10-cre mice were a gift from Dr. Chinfei Chen (Harvard Medical School). In the resulting Chx10-Cre;Ai32 mice, channelrhodopsin is expressed in the retinal ganglion cells and their axons, allowing us to optically stimulate them in brain slices (Litvina and Chen, 2017). Mice of both sexes were used for experiments and were raised in standard housing with standard feed and 12/12 h light/dark cycle.

#### Enucleation

We followed a previously described surgical procedure (Wilding et al., 2015) for bilateral enucleation with some minor modifications. 6-8 week-old mice were anesthetized with an intraperitoneal injection of ketamine (100 mg/kg) and xylazine (10 mg/kg). Once surgical level anesthesia was achieved (confirmed by the absence of toe-pinch and corneal reflexes), the eyes were swabbed with alternating swabs of betadine and 70% ethanol and one eye was enucleated with the help of fine ophthalmic surgical scissors. The orbit was flushed with sterile saline, followed by bupivacaine (0.5 %) and was packed with Gelfoam. The eyelid margins were then trimmed using fine ophthalmic scissors and sutured closed with two to three simple interrupted stitches. The procedure was repeated for the other eye. After the surgery, the mice were injected with Rimadyl (5 mg/kg) subcutaneously and allowed to recover.

#### Brain Slice Preparation

For electrophysiology experiments, we prepared acute brain slices at several time points after enucleation (2-4 days, 7-10 days, 14-16 days) for patch-clamp electrophysiology following the protected recovery method (Ting et al., 2014, 2018). Mice were euthanized with CO_2_ asphyxiation and cervical dislocation. The brain was immediately dissected and placed in a beaker with a frozen slurry of artificial cerebrospinal fluid (ACSF; 128.3 mM NaCl, 2.5 mM KCl, 1.25 mM NaH_2_PO_4_, 24 mM NaHCO_3_, 12.5 mM glucose, 2 mM CaCl_2_, 2 mM MgSO_4_) and allowed to chill for approximately 1 minute. The cerebellum was removed, and the brain was mounted onto the specimen holder. For parasagittal sections, the brain was cut with a razor blade angled at approx. 5 degrees from the longitudinal fissure and approx. 20 degrees angled laterally from the vertical axis after the protocol described by Turner and Salt (1998) and pieces were mounted cut side down on a block of agar in the specimen holder. We used a vibratome (Leica VT1000S) to cut 250 μm-thick coronal sections or 300 μm-thick parasagittal sections in ice-cold ACSF continuously bubbling with a mixture of 5% O_2_ and 95% CO_2_. We confirmed and selected slices with dLGN with the help of a dissecting microscope. Coronal slices were hemisected along the midline. Slices were transferred to a beaker containing N-Methyl-D-glucamine diatrizoate-ACSF (NMDG-ACSF; 107 mM NMDG, 2.5 mM KCl, 1.25 mM NaH_2_PO_4_, 30 mM NaHCO_3_, 20 mM HEPES, 0.5 mM CaCl_2_, 10 mM MgSO_4_, 2 mM thiourea, 5 mM L-ascorbic acid, and 3 mM Na-pyruvate) warmed to approximately 30° C and bubbled with 5% O_2_ and 95% CO_2_. After 10-15 minutes, we transferred and allowed the slices to recover for one hour in a beaker containing room temperature ACSF before beginning patch-clamp recording.

#### Electrophysiology

After one hour of recovery in ACSF, we transferred the slices to a recording chamber placed on an upright microscope fitted with an electrophysiology setup (Scientifica, Uckfield, UK or Olympus BX51WI). We targeted dLGN neurons under IR illumination for whole-cell patch-clamp recording using pipettes pulled from thin wall borosilicate capillary tubes with an internal filament. The pipettes (OD = 1.2 mm, ID = 0.9 mm) were pulled using a Sutter Instrument pipette puller (Sutter Instrument P-1000, Novato, CA) and had a resistance of 7-8 MΩ. Signals were visualized on a digital oscilloscope (Rigol DS1102E, Rigol USA, Beaverton, OR). Current clamp and voltage clamp recordings were obtained using Clampex 10.6 software (Molecular Devices, San Jose, CA), Multiclamp 700B amplifier (Molecular Devices) and a Digidata 1550B digitizer (Molecular Devices). Electrophysiological recordings were obtained at room temperature.

Spiking behavior and miniature excitatory post-synaptic currents (mEPSCs) were recorded from the dLGN of C57 mice. We used potassium-based pipette solution (in mM: 120 K-gluconate, 8 KCl, 2 EGTA, 10 HEPES, 5 ATP-Mg, 0.5 GTP-Na2, 5 phosphocreatine-Na2, 100 μM CF568 hydrazide dye (Sigma Aldrich, St. Louis, MO), and 1-2% Neurobiotin (Vector Labs, Burlingame, CA)) for these recordings. Reported voltages are corrected for liquid junction potential of 14 mV. mEPSCs were recorded in voltage-clamp configuration while holding the cells at -74 mV. TC neurons in the dLGN were identified as cells with oval somata and low voltage activated Ca^2+^ currents. Once mEPSCs were recorded, we switched to current-clamp configuration and recorded responses to hyperpolarizing (−20 and -40 pA) and depolarizing (+20 to +260 pA) current injections.

We calculated the mean frequency and amplitude for the first 100 mEPSCs detected in a 60s recording from each cell using MiniAnalysis (Synaptosoft, Fort Lee, NJ). Input resistance was calculated from the voltage response to a -20 pA current injection and resting membrane potential was obtained as an average measurement of baseline membrane potential in current-clamp without any stimulus.

For optogenetics experiments, we used Chx10-cre;Ai32 mice (channelrhodopsin expression in retinal ganglion cell (RGC) axons in the dLGN). All data were recorded in voltage clamp configuration and the intracellular solution used for the experiments was Cesium-based (120 mM Cs-methanesulfonate, 2 mM EGTA, 10 mM HEPES, 8 mM TEA-Cl, 5 mM ATP-Mg, 0.5 mM GTP-Na_2_, 5 mM phosphocreatine-Na_2_, 2 mM QX-314). Reported voltages for this intracellular solution are corrected for a 10 mV liquid junction potential. The ACSF was supplemented with 60 μM picrotoxin to isolate excitatory inputs from influences of feedforward and feedback inhibitory circuits and measure NMDA currents without the contamination from inhibitory inputs.

Once we broke into the cells, we recorded AMPAR (α-amino-3-hydroxy-5-methyl-4-isoxazolepropionic acid receptor)- and NMDAR (N-methyl-D-aspartate receptor)-mediated synaptic currents (EPSC_AMPA_ and EPSC_NMDA_) via optogenetic stimulation of RGC axons while holding the cells at -70 mV and +40 mV, respectively. Amplitude of peak current at -70 mV was the AMPAR-mediated EPSC (I_AMPA_) while the amplitude of current after 20 ms of onset of stimulus (to minimize contamination of I_NMDA_ with outward I_AMPA_) while holding the cell at +40 mV was used as the amplitude of NMDAR-mediated EPSC (I_NMDA_). Since single channel conductance of AMPARs has been shown to be a function of interaction between auxiliary subunit and AMPAR subunit, phosphorylation of the C-terminal domain, auxiliary and AMPAR subunit composition, and changes in the pore structure (Greger et al., 2017), changes in AMPA/NMDA ratio gives us a measure of changes in AMPAR number and composition while assuming that the NMDAR component remains constant (Chen and Regehr, 2000).

Paired pulse experiments were performed in the presence of bath-applied γ-DGG (*γ*-D-glutamylglycine, 200 μM) to prevent contributions from AMPAR saturation to short-term plasticity (Meyer et al., 2001) and CTZ (cyclothiazide, 100 μM) was included to minimize contributions from AMPAR desensitization (Trussell et al., 1993; Otis et al., 1996; Sakaba et al., 2002). Current responses to paired pulses of LED stimulus were recorded at varying interval between pulses (100 ms, 200 ms, 500 ms and 1000 ms pulse interval) while holding the cell at -70 mV. Paired pulse ratio was measured as the ratio of the peak of second EPSC to the peak of first EPSC (EPSC2/EPSC1).

For release probability (Pr) measurement, we utilized a high frequency LED stimulus (10 Hz pulse train). The cumulative amplitude of the responses to the 15^th^ to 30^th^ pulse fit with a straight line and extrapolated to the Y-axis. The ratio of the first EPSC to the Y-intercept of the fit provides Pr. The slope of the fit provides an estimation of the vesicle replenishment rate (Schneggenburger et al., 1999; Sakaba et al., 2002; Thanawala and Regehr, 2013, 2016; Neher, 2015).

For recording of excitatory cortical feedback synapses in parasagittal slices, a concentric bipolar stimulating electrode was positioned in the thalamic reticular nucleus and the ACSF was supplemented with 60 μM picrotoxin and 5 μM CGP55845 (Tocris). The corticothalamic axons were stimulated with pairs of current pulses separated by 50 ms (100 μA to 1 mA, 0.1-0.5 ms pulse duration).

TC neuron excitability was assessed by measuring the area under the curve (AUC) of the current-spike plot for individual cells and by the half-maximum current amplitude from a Boltzmann fit. Both analyses were implemented in GraphPad Prism.

#### Single cell fills

For single cell fills, we loaded the cells with neurobiotin (1%) and CF568 dye (100 μM) via passive diffusion during recording. Following electrophysiology, the slices were immediately fixed in 4% PFA for ∼1 day at 4° C. Following fixation, the slices were washed with PBS and incubated with streptavidin conjugated with Alexa fluor 568 (1 μg/ml) in permeabilization buffer containing PBS, 0.5% DMSO, and 1% Triton X-100 (pH adjusted to 7.4 with NaOH) for > 3 days at 4° C. The slices were then washed with PBS six times, mounted, and coverslipped on Superfrost plus slides with VectaShield hardset (Vectorlabs #H-1400-10) mounting medium.

#### Immunohistochemistry

For immunohistochemistry, C57 mice with appropriate treatment (control or enucleated) were euthanized with CO_2_ asphyxiation and cervical dislocation. Brains were immediately dissected, washed briefly in PBS and fixed in 4% PFA for 4 hours. Fixed brains were cryoprotected overnight in 30% sucrose and embedded in 3% agar. 50 μm coronal brain slices containing the dLGN were obtained using a vibratome, collected immediately on SuperFrost Plus slides and stored at -20 ºC. The slides were washed in PBS followed by blocking and permeabilization for 1 hour in blocking buffer (PBS, 0.5% Triton X-100, 5.5% donkey serum, 5.5% goat serum, pH ∼ 7.4). The slides were then incubated in primary antibodies diluted in blocking buffer (1:250 guinea pig-anti-vGlut2, Millipore AB255; 1:500 rabbit-anti-Iba1, Wako 019-19741) overnight – vGlut2 is a selective marker for retinogeniculate terminals in the dLGN (Fujiyama et al., 2003; Land et al., 2004; Yoshida et al., 2009; Koch et al., 2011; Seabrook et al., 2013), while Iba1 was used to label microglia. Following incubation with primary antibodies, the slices were washed in PBS, blocked for 1 hour and incubated with secondary antibodies diluted in blocking buffer (1:200, goat-anti-guinea pig IgG, Alexa Fluor 488, Invitrogen A11073; 1:200 donkey-anti-rabbit IgG, Alexa Fluor 568, Invitrogen A10042), washed and coverslipped with Vectashield Hardset mounting medium and stored at 4 ºC until imaged.

#### Imaging and Analysis

Single cell fills were imaged on a 2-photon setup (Scientifica) equipped with a Ti-Sapphire laser (Spectra-Physics, Santa Clara, CA) tuned to 800 nm. Four images per plane were acquired with 1 μm spacing between each plane. The dendritic arbors were traced in ImageJ (Schindelin et al., 2012) using the Simple Neurite Tracer plugin (Longair et al., 2011). Sholl analysis was performed using the Sholl analysis plugin (Ferreira et al., 2014) for ImageJ.

For vGlut2 imaging and analysis, images were acquired with the same setup as described above. Four images per plane were acquired with 1 μm spacing between each plane (Imaging parameters: laser power at ∼ 100 μW, scale: 2.077 px/μm). Grouped average projection was obtained for all the images with a group size of 4. Single optical sections from the middle of the Z-series were analyzed for vGlut2; brightness and contrast were adjusted identically for all images and images were converted to RGB. The RGB images were analyzed using the Synapse Counter plugin in ImageJ to detect and quantify the vGlut2 (parameters: particle size = 6-200 μm^2^), and the results were used to calculate the density per 1000 μm^2^, and the average size of vGlut2 puncta (μm^2^).

For analysis of microglia activation from Iba1-stained dLGN sections, we followed previously published approaches using a skeleton analysis to quantitatively analyze microglia morphology across entire images (Young & Morrison 2018; Morrison & Filosa 2013). Microglia were analyzed in a maximum intensity projection image of 40 planes (1 μm spacing) captured on the 2-photon microscope (370×370 μm, 2.077 px/μm). The number of microglia cell bodies were counted within the frame and the total process length and number of process endpoints per microglial cell was calculated for the entire image using the Analyze Skeleton ImageJ plug-in (Arganda-Carreras et al., 2010). A “process length threshold” of 1 μm was used to exclude skeleton fragments arising from the staining and image acquisition (Young & Morrison 2018).

#### Statistics

Unless otherwise noted, data are presented as mean ± SEM and sample sizes in terms of numbers of cells and mice are reported. To avoid pitfalls from pseudoreplication (Eisner, 2021), statistical significance was measured using one-way nested ANOVA with a Dunnett’s multiple comparisons post-hoc test. A nested t-test was used for pairwise comparisons. P<0.05 was considered statistically significant. GraphPad Prism, Microsoft Excel and Clampfit were used to perform statistical analyses.

## Results

To evaluate the effects of traumatic optic nerve injury in the mature dLGN, we bilaterally enucleated adult mice and probed for changes in physiology and structure of the ganglion cell axon terminals as well as TC neurons. Our goal was to determine the time-dependent effects of severe injury at the retinogeniculate synapse, the rate and the extent of synaptic dysfunction, and responses of postsynaptic TC neurons to a relatively sudden and drastic loss of synaptic partners. Our findings demonstrate that following such injury, presynaptic dysfunction proceeds in a rapid manner in the retinorecipient region of the thalamus and compensatory mechanisms initiate in response, altering the physiology of the postsynaptic TC neurons in the dLGN.

### Optic tract degeneration starts early following severe injury to the optic nerve

Our first goal was to determine the time course of RGC axon terminal degeneration in the dLGN following enucleation. To accomplish this, we obtained 50 μm thick histological sections of the dLGN at 2 days, 7 days, and 14 days post-enucleation and stained them for the retinal axon terminal marker vGlut2 (Figure 1). We utilized a semi-automated analysis to detect and quantify the presynaptic marker in images obtained from vGlut2 stained dLGN slices. Our analysis revealed that enucleation had a statistically significant effect on the density of vGlut2 puncta in the dLGN (p<0.0001, one-way nested ANOVA; control n = 7, 2-day n = 6, 7-day n = 6, 14-day n = 6 mice). Dunnett’s post-hoc multiple comparison tests revealed a statistically significant reduction in vGlut2 punctum density at 7d and 14d post-enucleation compared to control (p<0.0001). We did not detect a statistically significant difference in vGlut2 punctum density compared to controls at two days post-enucleation (p>0.05 Dunnett’s post-hoc test). Additionally, we did not detect any statistically significant change in the median vGlut2 punctum size following enucleation (p=0.31, one-way ANOVA). When we omitted the primary antibody and performed an identical analysis (n = 3 control dLGN sections), little signal was detected (detected punctum density = 0.17 + 0.12 puncta/1000 μm^2^), indicating that the remaining vGlut2 signal observed in the 7 and 14 day post-enucleation groups represents genuine signal rather than non-specific secondary antibody labeling.

**Figure 1.**
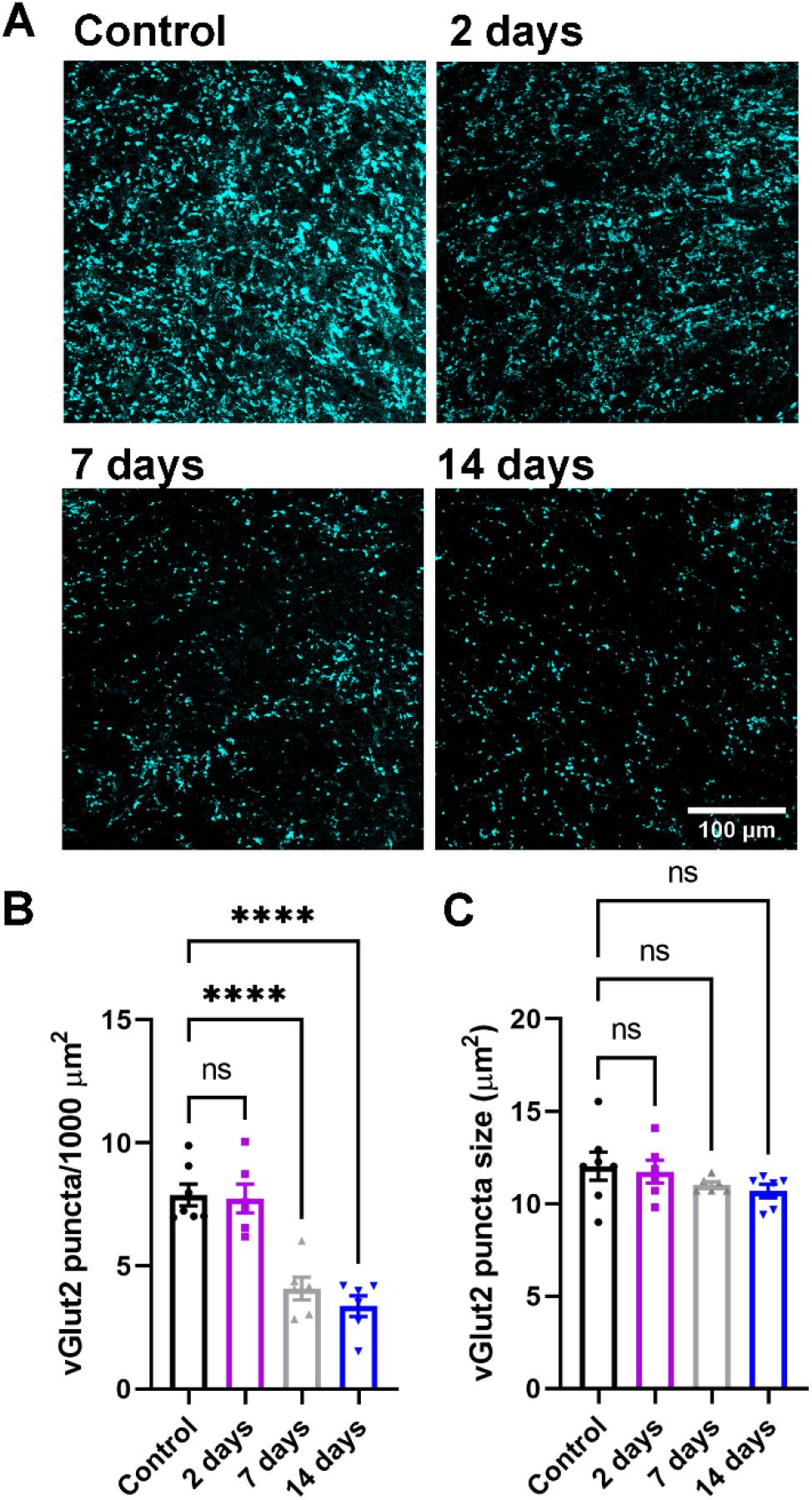
Loss of vGlut2-positive RGC terminals in the dLGN following enucleation. A) Single 2-photon optical sections of vGlut2 immunofluorescence staining from the dLGN following enucleation. B&C) Quantification of vGlut2 puncta density (B) and size (C). Each data point in C represents the median vGlut2 punctum size from an individual mouse. ****p<0.0001, Dunnett’s multiple comparison test. ns, not significant (p>0.05).

### Enucleation leads to microglial activation in the dLGN

Microglia are the resident immune cells of the central nervous system and are involved in synaptic pruning critical for refinement of retinogeniculate synapses. Moreover, in cases of disease or injury, activated microglia can mediate both protective and pathological functions, potentially recapitulating the developmental processes in adulthood (Hammond et al., 2018). Therefore, we probed whether enucleation led to microgliosis in the dLGN and, if so, over what time course this occurred (Figure 2). In control conditions (n = 7 mice), Iba1-stained microglia cell bodies were relatively sparse in the dLGN and morphology suggested they occupied a “resting/surveilling” state, with complex, fine processes radiating from the cell bodies. At 7 and 14 days post-enucleation (n = 6 mice for each group), the density of microglia in the dLGN was elevated compared to controls (Figure 2C) and quantification of the process endpoints and total branch length per cell revealed decreases in complexity (Figure 2D&E), consistent with an “activated” state. The timing of this activated morphology aligned with the timing of diminished vGlut2 staining (Fig 1).

**Figure 2.**
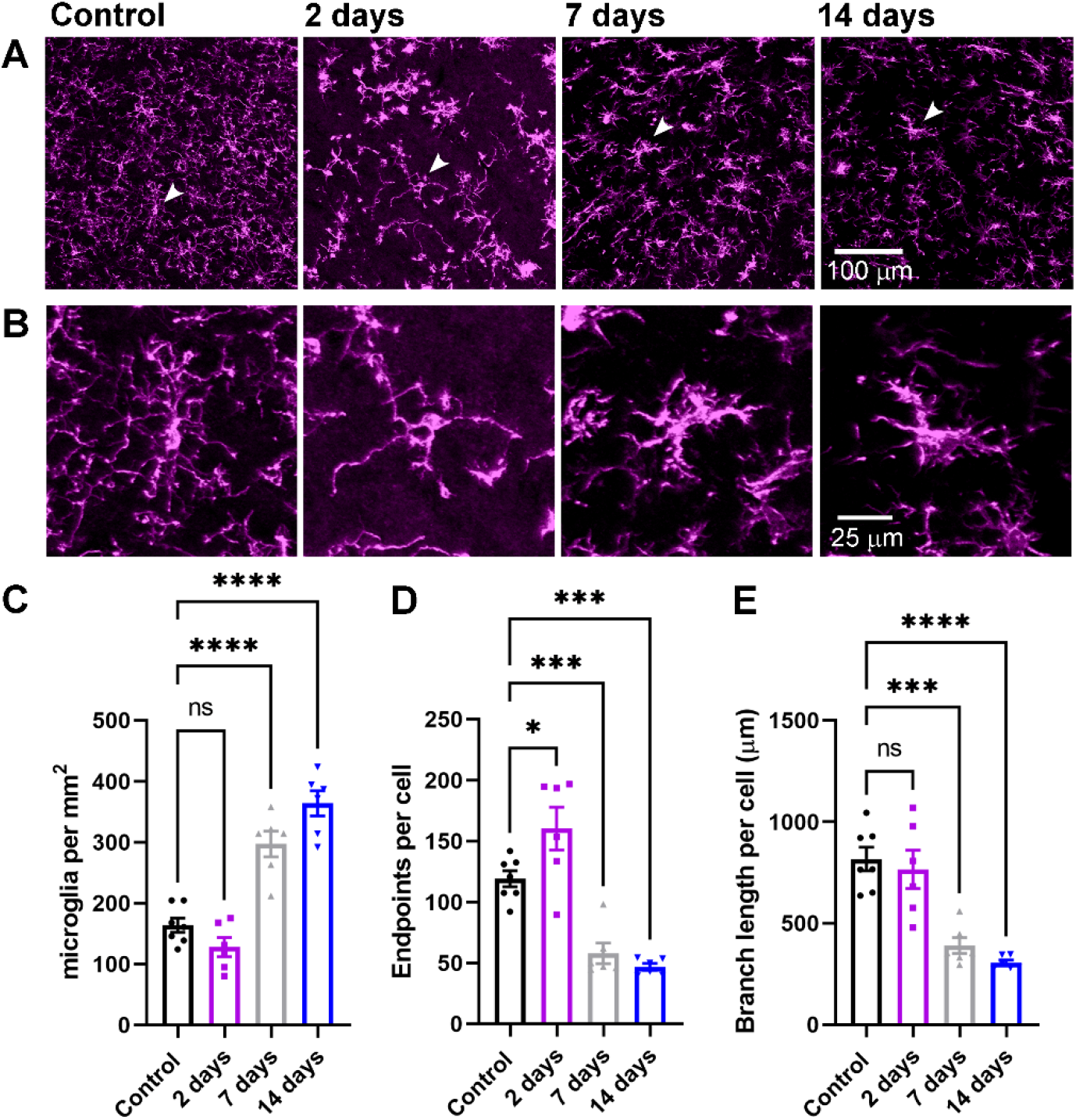
Evidence for microglia activation following bilateral enucleation. A) Maximum intensity projection of the dLGN stained with an anti-Iba1 antibody to label microglia. B) Higher magnification images of the individual microglia identified by arrows in the images in A. C) Quantification of microglia cell body density in the dLGN. D) Quantification of microglia endpoints per cell. E) Quantification of microglia branch length per cell. *p<0.05, ***p<0.005, ****p<0.0001, Dunnett’s multiple comparison test. Control n = 7 mice; 2 days n = 6 mice; 7 days n = 6 mice; 14 days n = 6 mice.

### Enucleation triggers progressive loss of retinogeniculate synaptic function

Having found that enucleation leads to a severe diminishment in vGlut2 labeling over the course of two weeks, our next set of experiments sought to determine how and over what time-course enucleation altered the function of synaptic inputs to dLGN TC neurons. To accomplish this, we measured evoked EPSCs from TC neurons using the Chx10-Cre;Ai32 mice in which channelrhodopsin is expressed in the retinal ganglion cells, allowing for activation of RGC axons in acute brain slices using 460 nm light (Figure 3). TC neurons were voltage-clamped at -70 mv to measure AMPA receptor-mediated EPSCs (EPSC_AMPA_) and at +40 mV to measure NMDA-mediated EPSCs (EPSC_NMDA_; Figure 2A). In these experiments, there was a statistically significant difference between groups for both EPSC_AMPA_ (Figure 3B) and EPSC_NMDA_ (Figure 3C) measurements (p<0.0001 and p=0.0004, respectively, nested ANOVA). EPSCs were detectable at 2-days post-enucleation but were reduced in amplitude relative to controls (p=0.007 for EPSC_AMPA_ and p=0.02 for EPSC_NMDA_, Dunnett’s post-hoc test) and were almost entirely absent by at 7d and 14d post-enucleation time points (p<0.005 for both, Dunnett’s post-hoc test). At 2d post-enucleation, when we could still measure detectable EPSC_AMPA_ and EPSC_NMDA_, the ratio of AMPA and NMDA EPSCs (AMPA/NMDA ratio, Figure 3D) was not significantly different between control and enucleation (p=0.92, nested t-test). These data indicate that retinogeniculate synaptic responses are lost over two weeks following enucleation, largely consistent with the diminishment of vGlut2 staining (Fig 1, above), but that there is no change in the number or composition of post-synaptic glutamate receptors at those synapses prior to loss of synaptic transmission (Greger et al., 2017; Chen & Regehr 2000).

**Figure 3.**
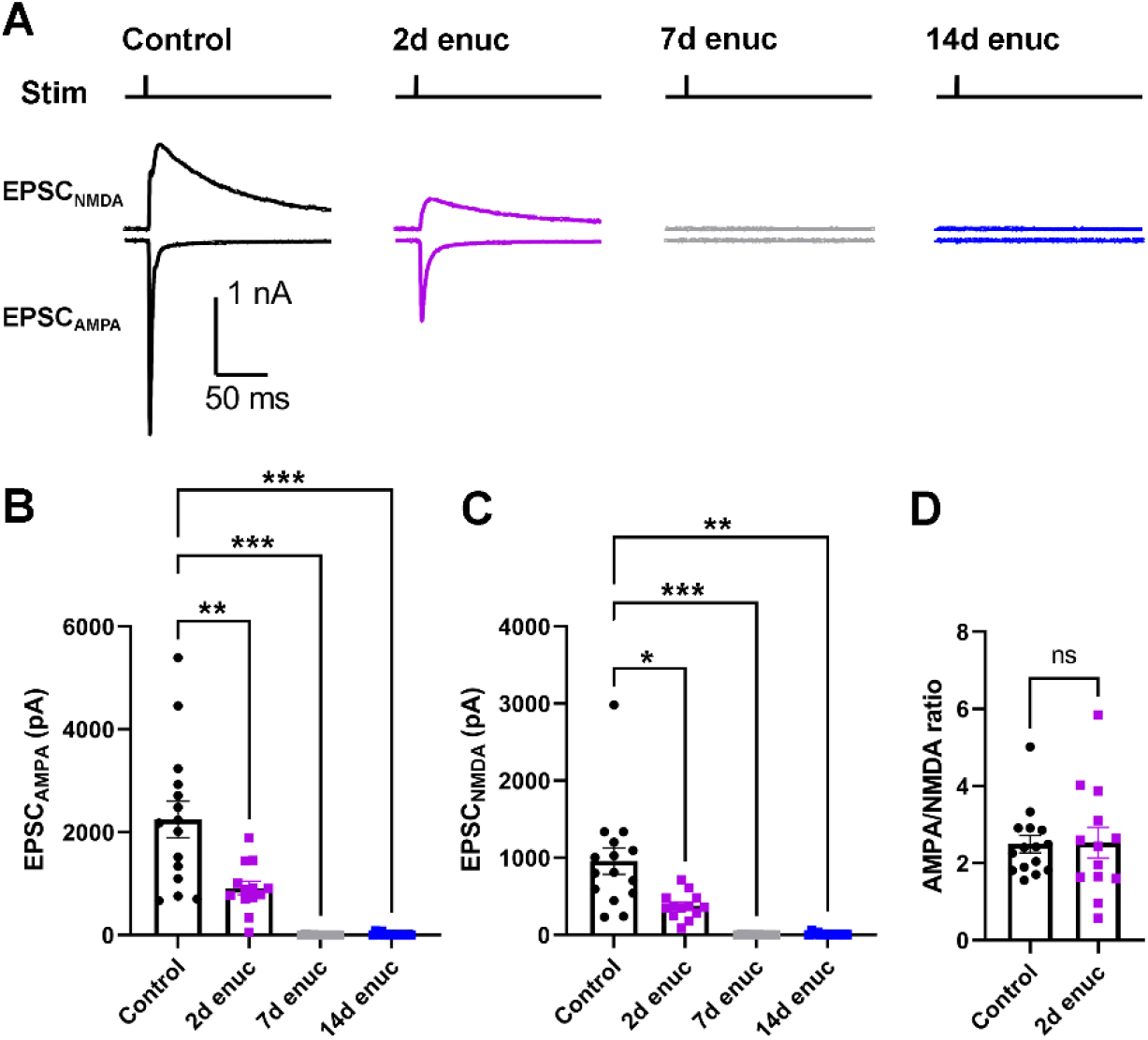
Loss of retinogeniculate synaptic function following enucleation. A) Representative traces of EPSCs recorded from the TC neurons in response to optogenetic stimulation of RGC axons from Chx10-Cre;Ai32 mice. AMPA receptor-mediated excitatory post-synaptic currents (EPSC_AMPA_) were recorded at -70 mV while NMDA receptor-mediated EPSCs (EPSC_NMDA_) were recorded at +40 mV. The stimulus timing is marked above the traces (“Stim”). B) Quantification of EPSC_AMPA_ amplitudes measured at the EPSC peak. C) Quantification of EPSC_NMDA_, measured 20 ms post-stimulus. Control n = 21 cells from 7 mice; 2d enuc n = 13 cells from 4 mice; 7 d enuc n = 10 cell 4 mice; 14d enuc n = 9 cells 3 mice. *p<0.05, **p<0.01, ***p<0.005, nested one-way ANOVA with Dunnett’s multiple comparison test. D) AMPA/NMDA ratio for control and 2-day post-enucleation recordings. (n.s., p>0.05, nested t-test).

In previous work from our lab, we demonstrated that ocular hypertension (OHT), which causes a more subtle optic nerve injury than the transection involved in enucleation, brings about an increase in retinogeniculate synaptic vesicle release probability (Pr) (Bhandari et al., 2019). Therefore, our next goal was to examine whether similar changes occur in retinogeniculate Pr in our model of severe injury to the optic nerve, after injury, but prior to total loss of synaptic responses or total degeneration of RGC axon terminals. Therefore, we tested for changes in Pr at 2-3 days post-enucleation. If this was the case, it might be a presynaptic attempt by retinogeniculate synapses to preserve synaptic strength following the injury associated with enucleation. We used two different approaches to estimate Pr. In our first set of experiments, we used dLGN sections of Chx10-Cre;Ai32 mice and evoked EPSCs with pairs of 460 nm LED stimuli separated by varying inter-pulse intervals (100 ms, 200 ms, 500 ms, and 1000 ms, Figure 4) in the presence of 100 μM cyclothiazide and 200 μM *γ*DGG to prevent AMPA receptor desensitization and saturation, respectively. The ratio of the second EPSC to the first EPSC (paired pulse ratio (PPR) = EPSC2/EPSC1) gives relative measure of Pr. We did not detect any significant change in the PPR at any interstimulus interval (p>0.05, nested t-test), suggesting that the presynaptic vesicle release probability was not significantly altered at 2 days post-enucleation (Figure 4B).

**Figure 4.**
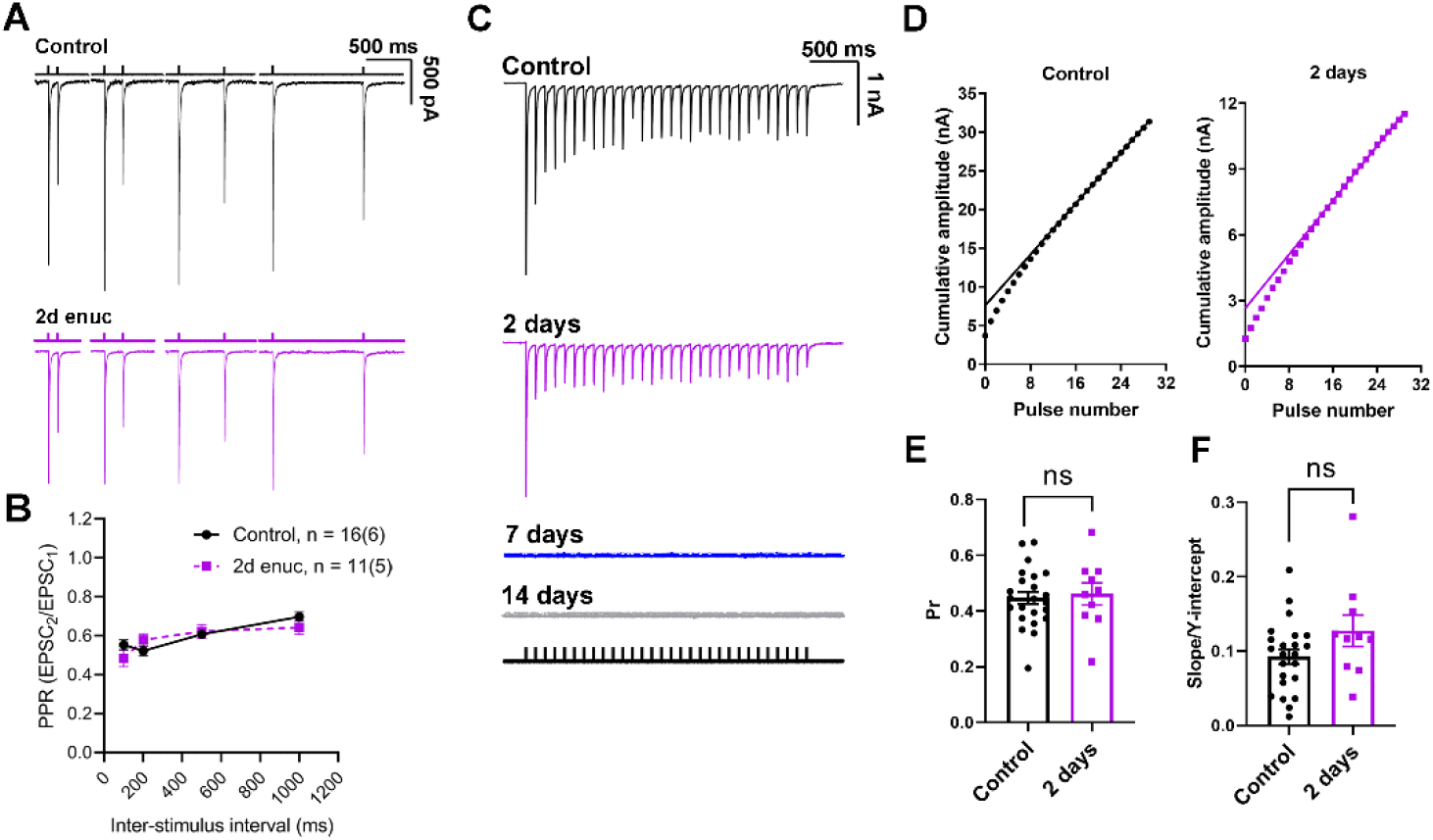
Presynaptic vesicle release probability at the retinogeniculate synapse does not change in response to bilateral enucleation. A) Example traces of EPSCs recorded in TC neurons in response to pairs of stimuli applied to stimulate the RGC axons spaced at different time intervals (100 ms, 200 ms, 500 ms, and 1000 ms). The bathing solution was supplemented with *γ*DGG (200 μM) and cyclothiazide (100 μM). B) Plot of paired pulse ratio at different stimulus intervals. Sample sizes are number of cells and (mice). There was no significant difference between control and enucleated at any time interval (nested t-test). **C)** Representative examples of excitatory post-synaptic current responses of TC neurons to a 10 Hz stimulus train at different time points following bilateral enucleation. The stimulus marker is shown below the traces. D) Cumulative EPSC amplitudes plotted against stimulus number with a linear fit of 15^th^ to 30^th^ data points extrapolated to the Y-axis. There was no detectable response at 7- or 14-day time points E) Quantification of release probability (Pr), measured as the ratio of the first EPSC to the Y-intercept. No significant difference was observed in the presynaptic vesicle release probability between the control (n = 23 cells from 7 mice) and the enucleated cohort at 2-days post enucleation (n = 10 cells from 4 mice, nested t-test). F) Slope of the linear fit, taken as a measure of vesicle pool replenishment rate and normalized to account for relative changes in EPSC size, was not significantly different between control and 2d post-enucleation (p>0.05, nested t-test).

In a parallel approach, we used a 10 Hz train of optogenetic stimuli to evoke EPSCs from TC neurons in the presence of cyclothiazide and *γ*DGG (Figure 4C-F). High-frequency stimulation induces a rapid release of vesicles from the presynaptic terminal leading to a state of equilibrium between the rate of vesicle replenishment and vesicle release. Subsequently, quantification of the amplitude of EPSCs evoked during high frequency stimulation and further analysis allows us to estimate Pr (Sakaba et al., 2002; Thanawala & Regehr, 2016). To obtain a measure of Pr at the retinogeniculate synapse, we plotted the cumulative EPSC amplitude and fit the 15^th^ to the 30^th^ responses with a straight line (Figure 5B). The ratio of the first EPSC amplitude to the Y-intercept of the fit is Pr. Comparable to the results obtained from our PPR experiments, we did not detect any change in Pr between control (0.45 ± 0.02; n = 23 cells, 7 mice) and enucleated animals at 2 days post injury (0.46 ± 0.04; n = 10 cells, 4 mice, p = 0.65, unpaired nested t-test; Figure 4E). Moreover, the rate of vesicle pool replenishment, measured as the slope of the linear fit normalized to the Y-intercept, was not altered 2 days post-enucleation (p = 0.12, unpaired nested t-test; Figure 4F). These results demonstrate that bilateral enucleation does not trigger any detectable change in the presynaptic vesicle release probability at retinogeniculate synapses within 2 days of injury.

**Figure 5.**
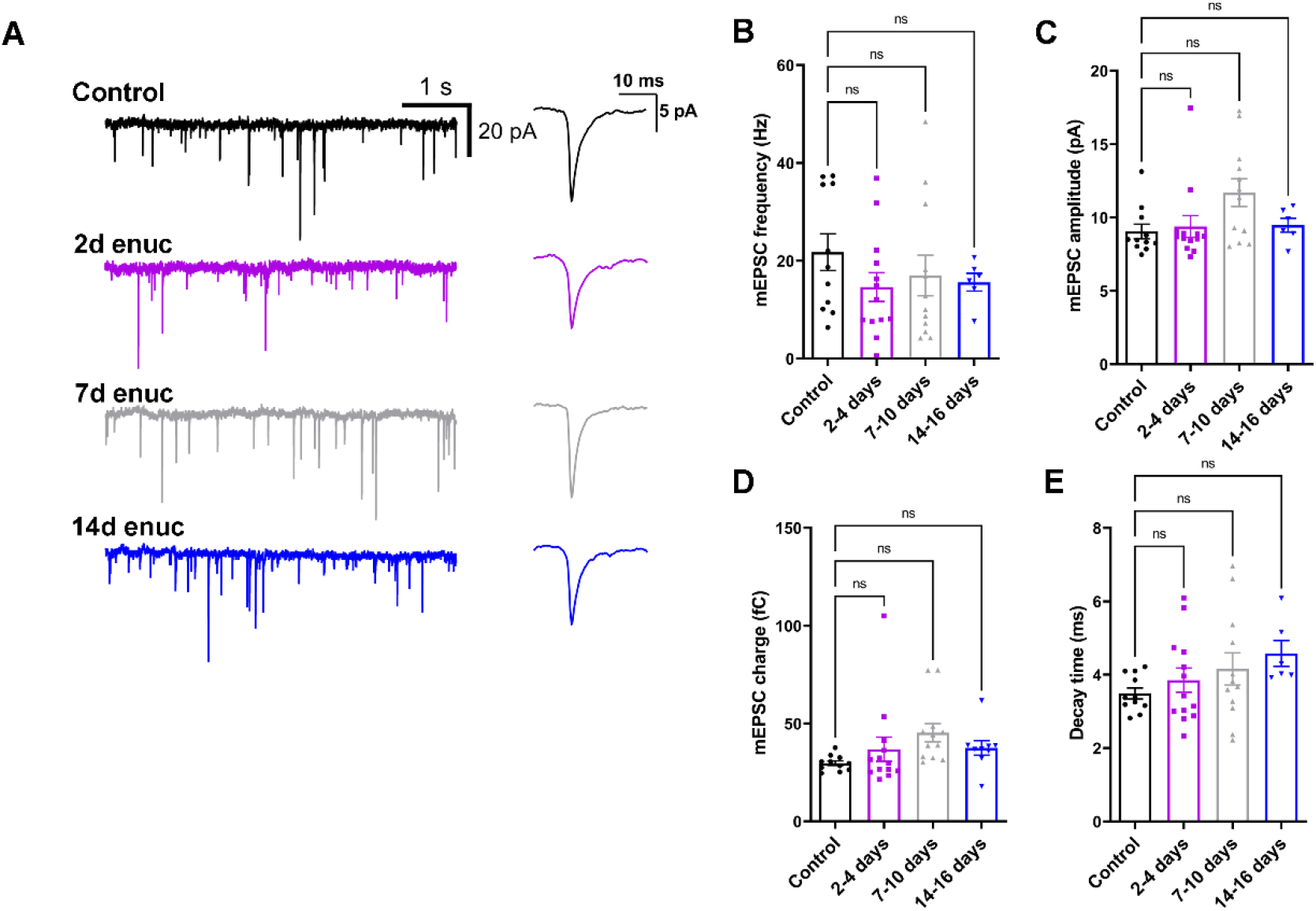
Bilateral enucleation does not alter single-vesicle post-synaptic current properties in the dLGN. A) Representative traces of mEPSCs recorded from dLGN TC neurons in control and enucleated animals in the absence of stimulation. Waveforms of average detected events are shown to the right. B) mEPSC frequency was not significantly altered following bilateral enucleation (nested ANOVA and Dunnett’s multiple comparison test). C) Quantification of mEPSC amplitude following enucleation. D) mEPSC charge was not significantly affected following enucleation. E) mEPSC decay kinetics were not significantly changed following enucleation. Control n = 11 cells from 5 mice; 2d enuc n = 13 cells from 8 mice; 7d enuc n = 12 cells 7 mice; 14d enuc n = 6 cells 4 mice.

Knowing that visual deprivation alters the balance of inputs arising from retinal and non-retinal sources in the dLGN (Krahe and Guido, 2011), and to test whether and how enucleation affects spontaneous excitatory drive to TC neurons, we next recorded miniature excitatory post synaptic currents (mEPSCs) from TC neurons of unoperated controls and enucleated animals (Figure 5). Surprisingly, we did not find a significant difference in mean mEPSC frequency at any time point following enucleation (nested ANOVA with Dunnett’s test, Figure 5B), despite the lower density of vGlut2-stained RGC synaptic terminals as shown in Figure 1, and loss of evoked synaptic response (Fig 3). Moreover, there was no significant difference in the mean mEPSC amplitude following enucleation (p>0.05; nested ANOVA with Dunnett’s test) compared to mEPSC amplitude from control animals. Additionally, we did not detect a statistically significant effect on mEPSC charge (Figure 5D).

Although retinogeniculate synapses are mostly lost by this time point, these results might point to a change in the properties of non-retinal excitatory inputs to TC neurons. Specifically, the lack of an effect of enucleation on mEPSCs might be a consequence of the fact that RGCs provide only ∼10% of the excitatory contacts onto TC neurons in the dLGN, with the remainder deriving largely from cortical sources. In this case, any changes in mEPSCs arising from RG terminals might be swamped by the comparatively greater number of non-retinal mEPSCs. Alternatively, loss of RGC inputs might be paralleled by a concurrent increase in the strength and/or number of non-retinal inputs, as appears to happen at TC neurons following visual deprivation (Krahe and Guido, 2011). Or, our result might be the result of a combination of those options. Corticothalamic inputs are concentrated at distal dendritic sites in TC neurons and slowing of the mean of the mEPSC decay kinetics following enucleation would be consistent with a greater relative proportion of events arising from non-retinal inputs. This is the consequence of dendritic filtering influencing mEPSC kinetics prior to detection by the patch electrode at the soma (Magee, 2000; Krahe & Guido, 2011). To test this possibility, we examined the mEPSC decay kinetics in slices from enucleated mice (Figure 5D). However, we did not find a statistically significant effect of enucleation on mEPSC decay kinetics in any of our experimental groups (p>0.05, nested ANOVA).

In a separate experiment, we tested whether enucleation leads to functional changes at cortical feedback synapses onto dLGN TC neurons. To accomplish this, we prepared parasagittal slices and stimulated corticothalamic (C-T) projections with an electrode positioned in the thalamic reticular nucleus. C-T excitatory synapses display pronounced paired pulse facilitation, a consequence of their low Pr (Turner & Salt 1998) and we tested for changes in Pr by stimulating C-T axons with pairs of stimuli separated by 50 ms (Figure 6) while measuring PPR of the evoked EPSCs (Figure 6A-B). Suggesting that enucleation did not lead to an increase in C-T Pr, the PPR was similar in slices from enucleated mice (8-9 days post-enucleation, 3.2+0.3, n = 12 cells from 4 mice) when compared to control mice (PPR = 3.5+0.3, n = 11 cells from 3 mice, p = 0.28, nested t-test). Additionally, suggesting enucleation did not affect the post-synaptic population of AMPA receptors, the AMPA/NMDA ratio at C-T synapses in enucleated mice (1.9+0.2, n = 11 cells from 4 mice) was not significantly different from controls (1.9+0.3, n = 6 cells from 3 mice, p = 0.74, nested t-test, Figure 6C-D). Thus, the function of C-T synapses in the dLGN appears to be unaffected by enucleation at this time point.

**Figure 6.**
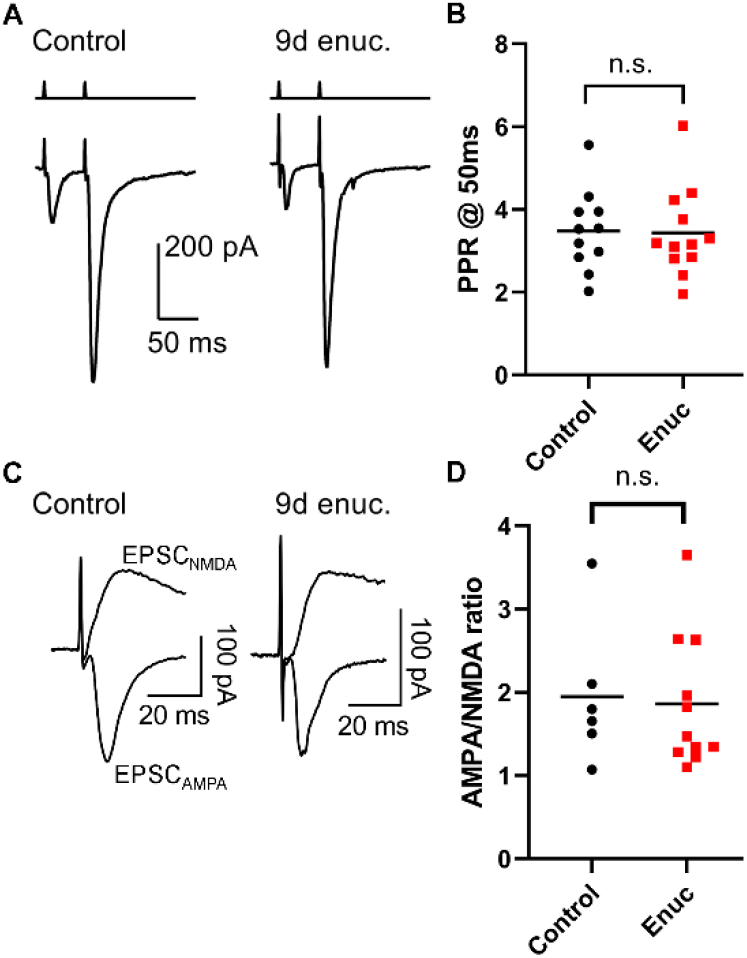
Bilateral enucleation does not alter short term plasticity or post-synaptic receptor complement at corticothalamic feedback synapses. A) Representative traces of excitatory post-synaptic currents (EPSCs) recorded in dLGN TC neurons in response to corticothalamic tract stimulation in parasagittal slices. Paired stimuli were separated by 50 ms. B) Quantification of paired pulse ratio (PPR) in recordings from control mice (n = 11 cells, 3 mice) and 8-9 days post-enucleation (n = 12 cell, 4 mice) shows that PPR was not significantly different between groups (p>0.05, nested t-test). C) Representative AMPA receptor- and NMDA receptor-mediated EPSCs recorded at -70 mV and +40 mV in response to corticothalamic tract stimulation. D) The AMPA/NMDA ratio at corticothalamic synapses was not significantly altered in mice at 8-9d post-enucleation (p>0.05, nested t-test).

### Bilateral enucleation increases the excitability of thalamocortical neurons in the dLGN

Having established loss of RG synaptic transmission within 7 days following enucleation, we next sought to investigate the effects on TC neuron spike output, which might reflect neuronal attempts at compensation in response to altered synaptic input. To accomplish this, we performed whole-cell current clamp recordings from TC neurons in 250 μm thick coronal brain slices from mice following enucleation. We utilized depolarizing and hyperpolarizing current injection steps to obtain spiking responses and measure TC neuron passive membrane properties. We observed that TC neurons in the dLGN from enucleated animals were more excitable, as evidenced by an increase in action potential firing in response to depolarizing current stimulation (Figure 7A-D). We quantified this by measuring both the area under the curve (AUC; nA*spikes) of the current-spike plots as well as the half-maximal current stimulus (I_50_) obtained from sigmoid fits. Following enucleation, the I_50_ was shifted leftward and a nested one-way ANOVA revealed a significant difference between groups (p=0.026) and post-hoc pairwise comparisons showed significant differences between control values and the 7-10 day (p=0.029) and >14d time points (p=0.041). Similarly, the AUC was increased at the >14-day time point compared to control (p=0.0076).

**Figure 7.**
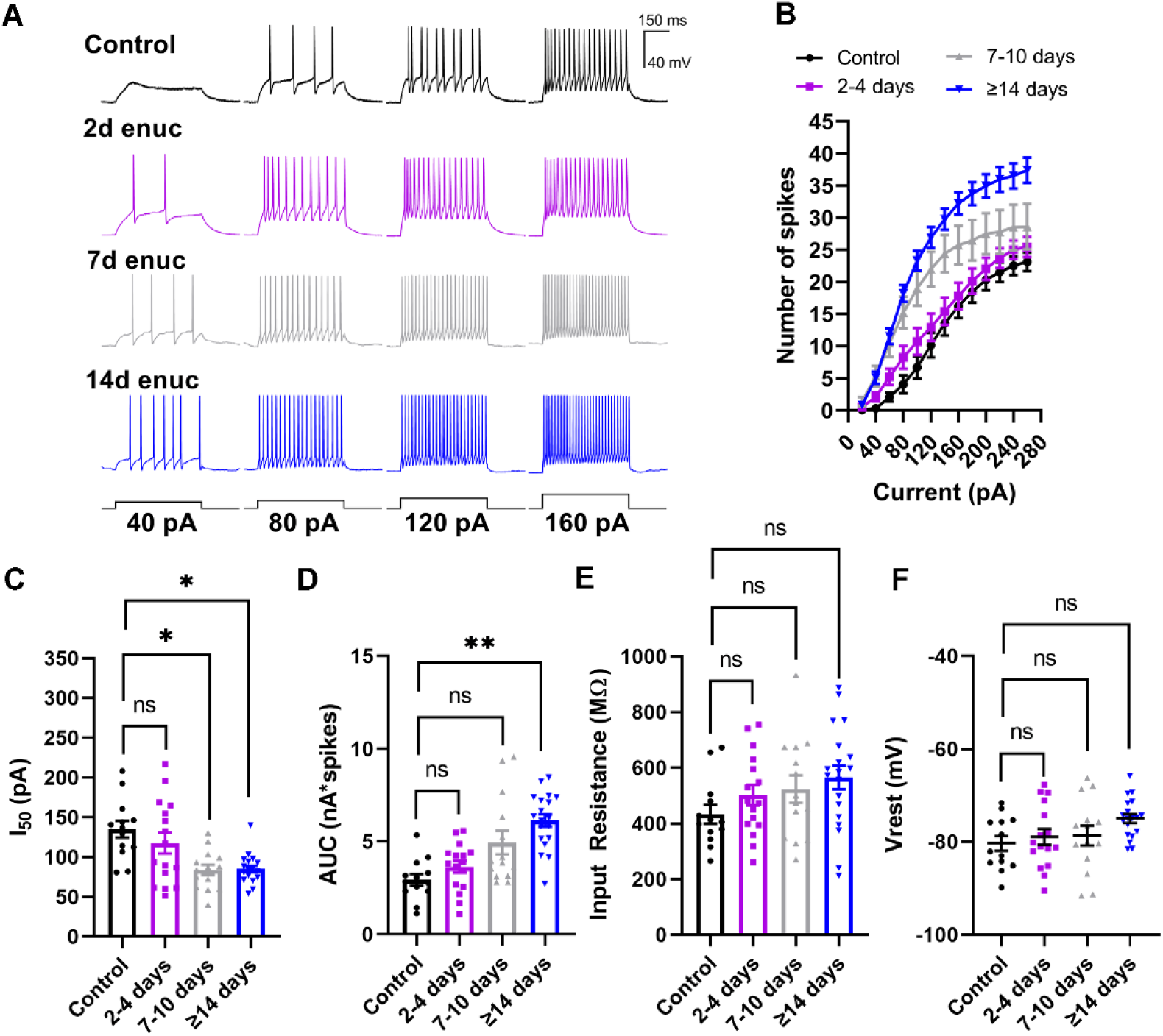
Bilateral enucleation increases the excitability of TC neurons. A) Whole-cell current-clamp recordings from TC neurons in response to 500-ms depolarizing current stimuli. B) Analysis of spiking data from TC neurons demonstrating enhanced excitability following enucleation. The number of evoked spikes were counted during the 500-ms stimulus and plotted against stimulus strength (Current). C and D) Analysis of the half-maximal current (I_50_) area under the curve (AUC) obtained from individual sigmoid fits. C) There was a significant reduction (leftward shift) in the I_50_ at 7-10-days and 14-16-days. D) There was significant increase in area under the curve at 7-10-days and 14-16-days post-enucleation. E) Input resistance, measured using the voltage deflection to a -20 pA current stimulus, was not significantly different between groups. F) Measurement of resting membrane potential (V_rest_) from the TC neurons showed no significant change in the V_rest_ between TC neurons from control animals and the enucleated cohort. *p<0.05, **p<0.01, Dunnett’s multiple comparison post-hoc test. Control n = 13 cells 7 mice; 2-4d enuc. n = 14 cells 7 mice; 7-10d enuc. n = 14 cells 8 mice; 14d enuc. n = 19 cells 9 mice.

We next sought to probe the origin of the increased excitability by analyzing the resting membrane potential (V_rest_) and the input resistance (R_in_). There was no significant difference in R_in_ between experimental groups (p=0.41, nested ANOA), and post-hoc pairwise comparisons did not show statistically significant differences between the control group and any time point post-enucleation. V_rest_ was similar in control and enucleated animals at all time points (Figure 7E, p=0.19, nested ANOVA) and post-hoc pairwise comparisons did not reveal significant differences between control and experimental groups. These results suggest that bilateral enucleation alters TC neuron spiking behavior, but not by any detectable alteration of TC neuron passive membrane properties.

### Altered dendritic structure in cells from enucleated animals

Optic nerve degeneration in glaucoma has been reported to alter the dendritic structure of dLGN relay neurons in primate and rodents (Gupta et al., 2007; Liu et al., 2014; Bhandari et al., 2019). Therefore, in a complementary set of experiments, we tested whether enucleation would trigger comparable post-synaptic dendritic changes. In this analysis, we examined single cell morphology and dendritic structure of TC neurons to evaluate enucleation-induced changes (Figure 8). The cells were filled with neurobiotin during electrophysiology recordings, processed with Alexa Fluor-conjugated streptavidin, and imaged later on a 2-photon imaging setup. Analysis of the dendritic tree using Sholl analysis pointed to a progressive pruning of TC neuron dendrites following enucleation (Figure 8A-B). There were no significant differences in dendritic structure between controls and enucleated animals at 2-4 days following enucleation. By 7-10 days, however, there was a subtle reduction in the number of dendritic intersections with the Sholl rings between 115-125 μm from the cell body. By 14 -16 days post-enucleation, dendritic fields were dramatically less complex. This reduction in dendritic complexity was more pronounced at the proximal portion of the dendritic structure relative to the soma. The reduction in dendritic complexity proximal to the soma could be represented by plotting the area under the curve (AUC) for the dendritic structure 35-105 μm away from the somata (Figure 8D). There also appeared to be a slight increase in dendritic complexity distal to the soma at 14 days post-enucleation. Analysis of the total dendritic length in the same set of images revealed no significant changes, (6555 ± 617.5 μm, n = 12 for controls, 7136 ± 1061 μm, n = 7, p = 0.646 for 2-4 days, 6144 ± 508.8, n = 5, p = 0.616 for 7-10 days, and 5618 ± 512.8, n = 9, p = 0.269 for 14-16 days; Figure 8E). This suggests that TC neuron dendrites are reorganized, with a delay, following loss of their presynaptic partners.

**Figure 8.**
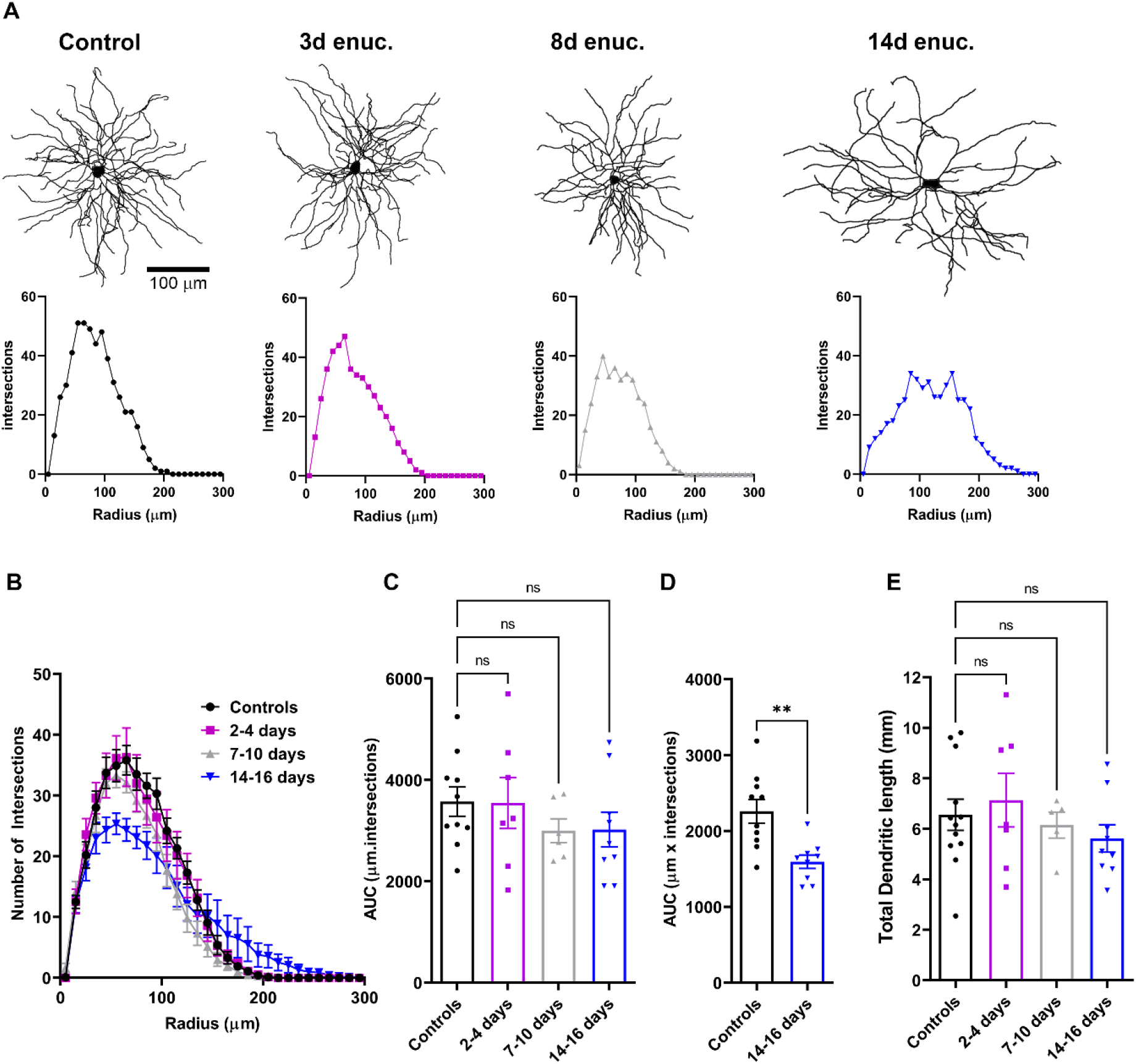
Reduction in dendritic complexity of TC neurons following bilateral enucleation. A) Examples of dendritic reconstructions of Neurobiotin-filled TC neurons (upper panels) from control animals and enucleated cohorts and corresponding individual Sholl analysis plots (lower panels). B) Sholl analysis plot showing the number of dendritic intersections with distance from the soma. Analysis of area under the curve (AUC) obtained from the Sholl plot did not reveal any statistically significant differences between the dendritic structure (C, n.s. p>0.05 Dunnett’s multiple comparison). However, when the area under the curve for the dendritic structure between 35-105 μm away from the TC neuron somata (D), there was a notable decrease in the dendritic complexity at 14-16 days compared to controls (**p<0.01, unpaired t-test). E) There was no significant change in the total dendritic length of TC neurons in all animals (n.s. p>0.05 Dunnett’s multiple comparison).

## Discussion

Neuronal activity is one of the most stable components of the nervous system and stability is required throughout life to maintain functional connectivity and activity. The balance in neuronal activity is the result of closely monitored processes that regulate network activity and adapt via compensatory mechanisms including change in synaptic properties, intrinsic excitability, and/or neuronal structure. Plasticity of synaptic transmission and in neuronal excitability in the visual cortex in response to altered visual input offers a prime example of this phenomenon (Turrigiano et al., 1998; Turrigiano, 1999; Desai et al., 2002; Turrigiano and Nelson, 2004; Wallace and Bear, 2004; Wierenga et al., 2005; Maffei and Turrigiano, 2008; Turrigiano, 2012; Keck et al., 2013; Nahmani and Turrigiano, 2014, Nys et al., 2015). Different experimental paradigms involving injury or visual deprivation have revealed that plasticity in the visual system extends to the thalamic partners as well (Bhandari et al., 2019; Krahe and Guido, 2011).

We had hypothesized that injury to the optic nerve and elimination of retinal input to the TC neurons triggers plasticity mechanisms that might represent neuronal attempts to compensate for the loss. Our results demonstrate that in response to bilateral enucleation and the consequent gradual degeneration of excitatory inputs from RGCs, dLGN TC neurons display both functional and structural plasticity, namely an increase in intrinsic excitability and a decrease in dendritic complexity. TC neurons from enucleated mice demonstrated an increased ability to fire action potentials in response to depolarizing current implying that their output to the visual cortex is enhanced in response to injury. The mechanisms underlying this are unclear and require further study; both resting membrane potential and input resistance were not significantly changed in TC neurons from control and enucleated mice, indicating that changes in spiking did not result from statistically detectable changes in proximity to spike threshold or increased membrane resistance. An alternative possibility is that enucleation and loss of synaptic inputs leads to changes in expression of TC neuron Na^+^ channels, similar to what occurs in retinal ganglion cells in glaucoma (Risner et al., 2018). Neuronal networks possess compensatory mechanisms that allow them to respond to changes in network activity (increasing or decreasing in input) either by scaling up or scaling down postsynaptic strength to maintain functional excitability and activity (Turrigiano, 2012; Chowdhury and Hell, 2018). Findings from our study demonstrate an enhancement of intrinsic excitability of TC neurons in response to loss of retinal inputs and is consistent with a scaling of neuronal responsiveness, although not synaptic function, following perturbation of synaptic inputs carrying visual information from the retina. Inhibitory interneurons in the dLGN also receive input from retinal sources and make feed-forward inhibitory synapses onto TC neurons. We did not explore functional changes to interneurons or to feed-forward inhibition onto TC neurons in this study, although those would be important areas for future studies.

The diminishment of vGlut2 staining in the dLGN by 7-14 days post-enucleation showed that RGC terminals progressively degenerate following transection of the optic nerve. This is further supported by the absence of EPSCs in TC neurons in response to optogenetic stimulation of RGC axon terminals by 7 days post-enucleation. Several studies have reported that in diseases like glaucoma, axonal degeneration and transport failure proceed in a distal to proximal manner and axonal pathology precedes RGC somatic degeneration (Crish et al., 2010; Calkins and Horner, 2012; Smith et al., 2016). Thus, the earliest pathological signs in diseases like glaucoma may not be axon or RGC soma loss, but functional changes in neuronal communication. This has been observed in glaucoma models, where the density of retinal terminals in the brain remains largely unchanged until late in the disease process, but they show a reduction in the size of mitochondria, reduction in trace metal levels, and reduction in the number and size of pre-synaptic active zones, suggesting physiological and metabolic deficiency and altered synaptic signaling (Crish et al., 2010; Smith et al., 2016). Moreover, microglial phagocytosis of RGC axon terminals is important in developmental refinement of retinogeniculate synapses (Schafer et al., 2012) and re-activation of microglia in adulthood in response to disease or injury might re-awaken those developmental processes and contribute to pathology (Hammond et al., 2018; Stevens and Schafer 2018). We found that microglia density in the dLGN increases with enucleation and those microglia display morphological features indicative of activation, including reduction in process length and number of process endpoints. There is evidence for microglia activation in the dLGN in experimental glaucoma and inflammatory demyelinating disease (Tribble et al., 2021; Fujishiro et al., 2020; Lam et al., 2009; Araujo et al., 2017). Although enucleation is a much more rapid and extreme axonal injury than typical glaucoma, our findings might recapitulate some of the pathological events of glaucoma or other optic neuropathies, albeit on an accelerated time course. Loss or dysfunction in synaptic connectivity in such cases can result in similar synaptic dysfunction and understanding response mechanisms could provide valuable insights to understanding disease pathology.

Synaptic scaling via changes in post-synaptic receptor complement is common in homeostatic plasticity, including in cortical pyramidal neurons (Maffei and Turrigiano, 2008; Turrigiano, 2008; Gainey et al., 2009; Keck et al., 2013; Fernandes and Carvalho, 2016; Teichert et al., 2017; Chowdhury and Hell, 2018a; Venkatesan et al., 2020). However, we did not observe any change in the amplitude of mEPSCs or I_AMPA_/I_NMDA_ indicating that there were no detectable changes in the number and/or composition of post-synaptic AMPARs at two days post-enucleation. We were unable to measure AMPA/NMDA at later time points due to the loss of retinogeniculate synaptic responses. Future studies should perform staining for AMPA and NMDA receptors to determine whether and over what time course postsynaptic sites are reorganized relative to presynaptic degeneration. Changes in AMPA/NMDA ratio in the dLGN have been previously reported during visual deprivation and dark rearing during development (Hooks and Chen, 2006; Louros et al., 2014). We did observe parallel reductions in the amplitude of EPSC_NMDA_ and EPSC_AMPAR_ after 2 days of enucleation. This suggests that the synapse is degenerating at that time point and there is reduced synaptic input to evoke stronger EPSCs. Alternately, as noted above, physiological, and metabolic deficiency in the optic nerve is present in a mouse glaucoma model and are accompanied by changes in RGC presynaptic active zones in the superior colliculus. If this were the case in the dLGN following enucleation, this could offer us an alternate explanation to the observed reduction in EPSC amplitudes. Notably, evoked EPSCs were entirely absent in experiments performed at 7d post-enucleation despite the persistence of some vGlut2-positive structures in the dLGN, suggesting that synaptic function might be lost prior to total loss of RGC axon terminals. Additionally, dLGN TC neurons appear to have a diverse array post-synaptic dendritic structures, with some retinal inputs occurring at spines and others onto dendritic shafts (Wilson et al., 1984). While probing for changes in dendritic spine morphology and/or number was beyond the scope of this study, it would be an interesting avenue to explore whether, and along what time course, loss of presynaptic inputs leads to structural changes at post-synaptic structures.

One of the most surprising results in this study is that we did not detect any statistically significant change in mEPSC frequency when we compare the means between different treatments although we observed prominent degeneration of RGC axon terminals. Furthermore, as Pr at the retinogeniculate synapse was unaltered at 2 days post-enucleation and as the cortical synapses vastly outnumber retinal contacts to a TC neuron (Connelly et al., 2016), these results represent the possibility that most of the mEPSCs arise from non-retinal inputs and retinal inputs make a minor contribution to mEPSCs recorded from the TC neurons in the dLGN. Indeed, ∼90% of excitatory inputs to TC neurons arise from corticothalamic feedback, although those synapses are considerably weaker than the less numerous retinogeniculate inputs. The variability in mEPSC frequency in TC neurons might make detection of lower numbers of retinogeniculate mEPSCs challenging, especially if mEPSCs are dominated by those arising from corticothalamic feedback synapses. Results from vGlut2 staining showing a diminished density of RGC axon terminals taken together with the lack of effect on mEPSCs supports the hypothesis that the majority of mEPSCs potentially arise from non-retinal synaptic inputs. In contrast to a prior study showing that altered visual input resulting from monocular deprivation leads to an increase in Pr from C-T excitatory inputs (Krahe and Guido, 2011), we did not detect any change in C-T synaptic function following enucleation. It remains possible that C-T synapses are altered at later time points, however.

Strikingly, we found that enucleation was followed by a loss of TC neuron dendritic complexity, especially in regions proximal to the cell body. One interesting feature of dLGN TC neurons synaptic anatomy is that inputs arising from the visual cortex are concentrated towards the dendrites distal to the somata while the synaptic inputs from the retina are concentrated towards the proximal dendrites (Sherman and Guillery, 2002; Krahe and Guido, 2011; Wilson et al., 1984; Connelly et al., 2016). This might suggest that the loss of synaptic input from the retina can lead to measurable changes in the dendritic structure. Changes in the dendritic component are fairly well-documented and occur during development and depend on retinal input (Vaughn et al., 1988; McAllister, 2000; Cohen-Cory and Lom, 2004; Cline and Haas, 2008; Callaway and Borrell, 2011; El-Danaf et al., 2015). Additionally, we have shown that in ocular hypertension, dLGN TC neuron dendritic complexity is reduced, especially in regions proximal to the TC neuron somata, where they receive retinal inputs (Bhandari et al., 2019). Findings from the current experiments indicate that similarly, there is a reduction in dendritic complexity proximal to the TC neuron somata, and these changes are prominent and significant by 14 days post-enucleation. Interestingly, these changes occur with a delay following diminished vGlut2 staining and loss of retinogeniculate synaptic function. This reduction in dendritic largely seems to match the reported sits retinogeniculate synaptic contact (30 -100 μm away from the somata), suggesting retinorecipient dendritic sites are lost while the more distal dendrites, which receive cortical inputs and are still synaptically active, remain intact. These results also support the notion that dendritic pruning and reorganization, even by neurons that are not directly damaged, is a common feature of degenerative disease and nervous system injury.

In conclusion, we found that enucleation leads to structural and functional changes in the dLGN. On the pre-synaptic side, these include loss of RGC presynaptic terminals and synaptic input to the TC neurons without any detectable changes in vesicle release probability. Post-synaptically, we observed an increase in the intrinsic excitability of TC neurons while no change in the post-synaptic glutamate receptor composition and number was observed prior to loss of synaptic function. This implies that TC neurons display certain features of homeostatic plasticity in response to degeneration of their major driver input and sheds light on mechanisms that might contribute to visual system plasticity in visual system diseases such as glaucoma or other optic neuropathies. This is important as plasticity mechanisms exist as stabilizing tools to modulate neuronal/synaptic function in response to any changes in the normal activity (increase or decrease), and while it has been extensively studied in the developing visual system (Gordon and Stryker, 1996; Desai et al., 2002; Turrigiano and Nelson, 2004; Hooks and Chen, 2006, 2008; Karmarkar and Dan, 2006; Turrigiano, 2008; Keck et al., 2013; Louros et al., 2014; Nahmani and Turrigiano, 2014) less is known about plasticity mechanisms operating in the adult. Our results provide an important groundwork on plasticity in the mature dLGN in response to optic nerve injury.

## Acknowledgements

We would like to thank Elizabeth Bierlein for comments on the manuscript. We are grateful to Dr. Chinfei Chen (Harvard Medical School) for supplying Chx10-Cre mice.

## Author contributions

AB and MVH designed the experiments; AB, JS, TW, and MVH performed the experiments; AB, JS, TW, and MVH analyzed the data; AB and MVH wrote the manuscript and prepared figures; MVH acquired funding.

